# Physiological bias governs neutrophil inflammatory threat perception

**DOI:** 10.1101/2024.04.05.588243

**Authors:** Denise Pajonczyk, Lina Pauli, Charlotte Pünt, Merle F. Sternschulte, Olesja Fehler, Thomas Vogl, Oliver Soehnlein, Marcel Bermudez, Carsten Alexander Raabe, Ursula Rescher

## Abstract

**Background and Purpose:** The functional G protein-coupled receptor (GPCR) signalling unit consists of an agonist acting on a receptor that is coupled to a G protein that transduces the signals to effectors within a complex cellular environment. While much attention is given to GPCR-agonist or GPCR-transducer relationships, the contribution of the cellular environment remains significantly unexplored.

**Experimental Approach:** Here, we juxtaposed the signalling responses triggered by the activation of two GPCR pattern recognition receptors, Formyl peptide receptor 1 and Formyl peptide receptor 2, in a recombinant cell system against their signalling dynamics in the native neutrophilic environment.

**Key results:** We observed that agonist activation leads to cell context-dependent substantial differences in the receptor signalling texture. While the impact of receptor activation on *de novo* cAMP formation varied depending on the cell type, MAPK activation was similar in both systems. This physiological bias was conserved across species. Expression analysis unveiled the absence of the Gαi-sensitive adenylyl cyclases ADCY5 and ADCY6 in neutrophils, implying that cAMP *de novo* synthesis cannot be inhibited by Gαi-coupled receptors. The signalling behaviour of the Gαi-coupled LTB4 high-affinity receptor BLT1 in neutrophils corroborated our findings.

**Conclusion and Implications:** Our data underscore the profound impact of the specific cellular environment on GPCR signalling, causing physiological bias in GPCR signalling, thereby affecting drug efficacy and therapeutic targeting.

## Introduction

Neutrophils, which represent 50–70% of the total white blood cell count, counterattack pathogenic threats through directed migration toward the site of infection and subsequent phagocytic clearance [1]. This critical immune cell population enhances the innate immune response by releasing pro-inflammatory mediators such as leukotriene B4 (LTB4) [2]. Key receptors on the neutrophil surface that detect stimulatory and activating signals are G protein-coupled receptors (GPCRs), including the β1- and β2-adrenoceptors [3], the high-affinity LTB4 receptor BLT1 [4], and the formyl peptide receptors FPR1 and FPR2 [5,6] which alert neutrophils to bacterial infection or tissue damage [7].

Downstream signalling of GPCRs includes adenylyl cyclases (ADCY) as effector proteins [8]. ADCYs catalyze the formation of cyclic adenosine monophosphate (cAMP) from ATP and are key regulators in the GPCR signalling pathways that rely on cAMP as the second messenger [9]. Through binding to cellular targets, cAMP regulates various physiological processes, including cell proliferation, metabolism, and apoptosis. ADCY isoforms typically synthesize cAMP upon activation via a stimulatory G protein alpha subunit (Gαs) of the heterotrimeric G protein complex. They can also be activated by forskolin (FSK), a plant diterpene that directly binds to the ADCY active site [10,11]. ADCY5 and ADCY6, which are structurally closely related, are activated by Gαs and FSK and inhibited by the inhibitory G protein alpha subunit (Gαi), thereby suppressing cAMP synthesis [12].

Observations of agonist-dependent differences in signalling capacities and the agonist-dependent preferred activation of one signalling pathway over other responses at the same GPCR have led to the concept of ‘biased signalling’ [13–17]. This concept has considerably expanded the initial view of a GPCR switching between a defined on-and-off state. Selective pathway activation out of the entire receptor signalling repertoire is now being exploited for developing novel drugs tailored to regulate disease-relevant receptor-linked pathways. In addition to ligand bias, a receptor’s response is also determined by cellular factors such as downstream transducers and effectors [17,18]. This confinement of agonist-driven signalling within the premises of the actual cellular environment is conceptualized as system bias [17]. However, this holistic view on the interplay of agonist/receptor/environment is still emerging. Due to ease of maintenance, heterologous expression systems, including recombinant GPCR-expressing HeLa and HEK293 cell lines, are common tools in GPCR research [19,20]. However, the results obtained in these recombinant models may not translate into the desired physiological or therapeutic effect *in vivo* due to unaccounted influences of system bias.

Since the FPRs and BLT1, like most chemoattractant receptors [21], are predominantly associated with Gi proteins [5,6], we focused on two primary G protein-mediated signalling pathways: (i) the Gαi-dependent inhibition of *de novo* cAMP accumulation and (ii) phosphorylation of the extracellular signal-regulated kinase 1/2 (ERK1/2) [22,23]. The unambiguous interpretation of our results was ensured by the use of receptor-specific ligands that activate only the designated GPCRs. From the vast spectrum of FPR ligands, we utilized only peptide agonists, as FPR activation through lipid agonists remains debated [16,24].

Our findings imply that the functional outcome of FPR1 and FPR2 activation is strongly determined by the cellular environment. Both receptors coupled to Gαi in both systems, however, agonist-driven inhibition of *de novo* cAMP formation, whether by direct ADCY activation through FSK or through activation of the Gαs-coupled β-adrenoceptors, was detected only in the heterologous context. Notably, FPR-mediated ERK1/2 activation was observed in both systems. This signalling texture was recapitulated in murine neutrophils and also by the LTB4 receptor BLT1. On the molecular level, the absence of Gαi-regulated ADCY activity correlated with the absence of ADCY5 and ADCY6, indicating that in neutrophils, Gαi-coupled receptors are incapable of counteracting a rise in cAMP levels. Thus, a generic signal transmitted by the same receptor leads to discrete responses, dependent on the specific identity of the receiving cell.

## Results

### Gai-dependent inhibition of cAMP *de novo* generation is cell system-dependent

To elucidate the impact of the cell environment, we compared the FPR-mediated signalling in HEK293 cells (negative for expression of both FPR1 and FPR2) ectopically expressing either FPR1 or FPR2 [25] to DMSO-differentiated HL60 (dHL60) cells that mimic neutrophils [26,27].

We utilized a FRET-based assay to assess receptor-induced changes in the levels of cAMP, the classical second messenger. In these cell systems, *de novo* cAMP generation could be robustly elicited through treatment with the direct adenylyl cyclase activator forskolin, albeit to a different extent (Suppl. Fig. S1B).

As expected, the pan-FPR agonist WKYMVm performed as a highly potent agonist in HEK293 FPR1 and HEK293 FPR2 cells. The FPR1 agonist fMLF favoured FPR1 over FPR2, and the FPR2-specific agonist MMK 1 [28,29] was selectively active at FPR2. Data agreed with a four-parameter logistic (LL4) model with the Hill slope set to 1. Surprisingly, no reliable responses were detected in agonist-activated dHL60 cells (Fig.1A) and the LL4 model was not applicable. Statistical analysis confirmed the different behaviour of HEK293-FPR and dHL60 cell lines (Fig. 1B). The unexpected outcome seen in dHL60 cells prompted us to test for a stimulatory activity on cAMP formation upon FPR activation. No changes in cAMP levels were observed when HEK293 FPR1 and HEK293 FPR2 cells were challenged with 10^-5^ M of each of the agonists and only minor increases were recorded in dHL60 cells (Fig. 1C). Also, these cells did not respond over a range of agonist concentrations, thus ruling out a bell-shaped-response (Suppl. Fig. 1). FPR-mediated inhibition of cAMP production was almost completely abolished in FPR-expressing HEK293 cell lines challenged with 10^-5^ M of each of the agonists in the presence of the Gi/o inhibitor pertussis toxin (PTX) [30], confirming the Gi-coupling of FPR1 and FPR2 in these cells (Fig. 1D). In contrast, PTX-treated dHL60 again failed to respond to agonist treatment, arguing against the capability of the receptors to drive cAMP formation when Gαi was inactivated (Fig. 1D).

**Figure 1.**
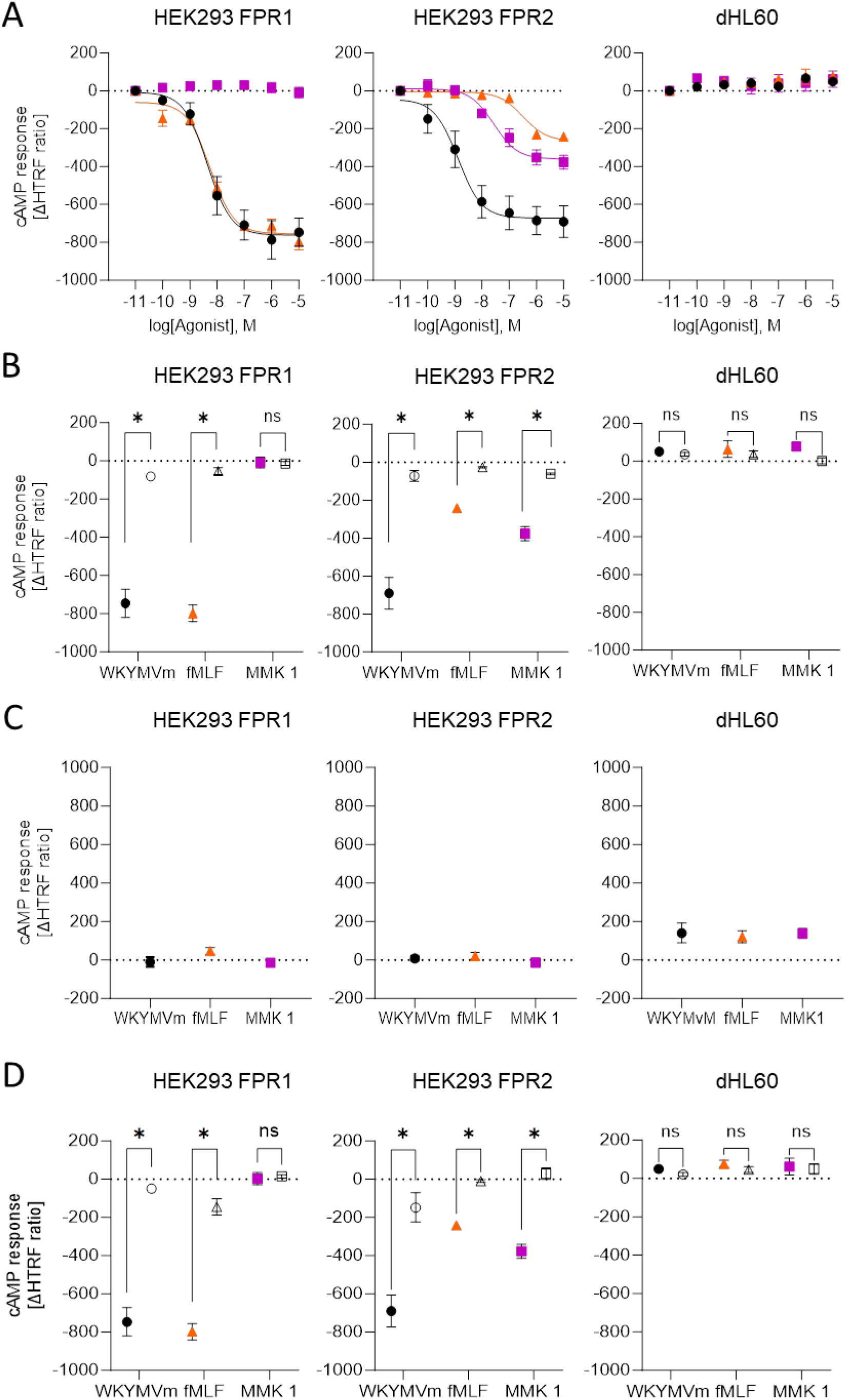
FPR-mediated effects on cAMP production depend on the cellular context. (A) The transgenic HEK293 FPR and the neutrophil-like dHL60 cell lines were stimulated with forskolin and the pan-FPR agonist WKYMVm, (black circle), FPR1-favoring fMLF (orange triangle), and the FPR2-specific agonist MMK 1 (magenta rectangle). ΔHTRF ratios were recorded 30 min after agonist addition. Data points represent the mean ± SEM ΔHTRF ratios of at least 5 independent experiments. Concentration-response curves were fitted using a four-parameter logistic model with the Hill slope set to 1. For data on agonist-mediated cAMP changes in dHL60, this mathematical model was not applicable. (B) Statistical analysis of the differences in responses at the lowest and the highest agonist concentrations. (C) Cells were evaluated for FPR-mediated changes in cAMP levels in the absence of FSK. Cells were treated with 10^-5^ M of the indicated agonists. (D) Cells pre-incubated with or without PTX were stimulated with 10^-5^ M each of the agonists and forskolin. Data points represent the mean ΔHTRF ratios ± SEM of at least 5 independent experiments. Mean ΔHTRF ratios ± SEM were compared by two-tailed Student’s *t*-test, ns, not significant, *, p-value < 0.05.

To rule out that the endogenous FPRs in dHL60 cells were not able to counteract the forced direct ADCY activation by forskolin, we next tested the agonists’ capacities to counteract cAMP production induced by the activation of a Gas-coupled receptor. We chose the β-adrenoceptor agonist isoproterenol [31]. Notably, the response exceeded the FSK-induced cAMP production and reached comparable levels in both cell models. (Suppl. Fig. S1). The FPR agonists fMLF and MMK 1 suppressed the isoproterenol-induced *de novo* cAMP production in a concentration-dependent manner in the FPR-expressing HEK293 cells only (Fig. 2), in line with the results observed in FSK-stimulated cells.

**Figure 2.**
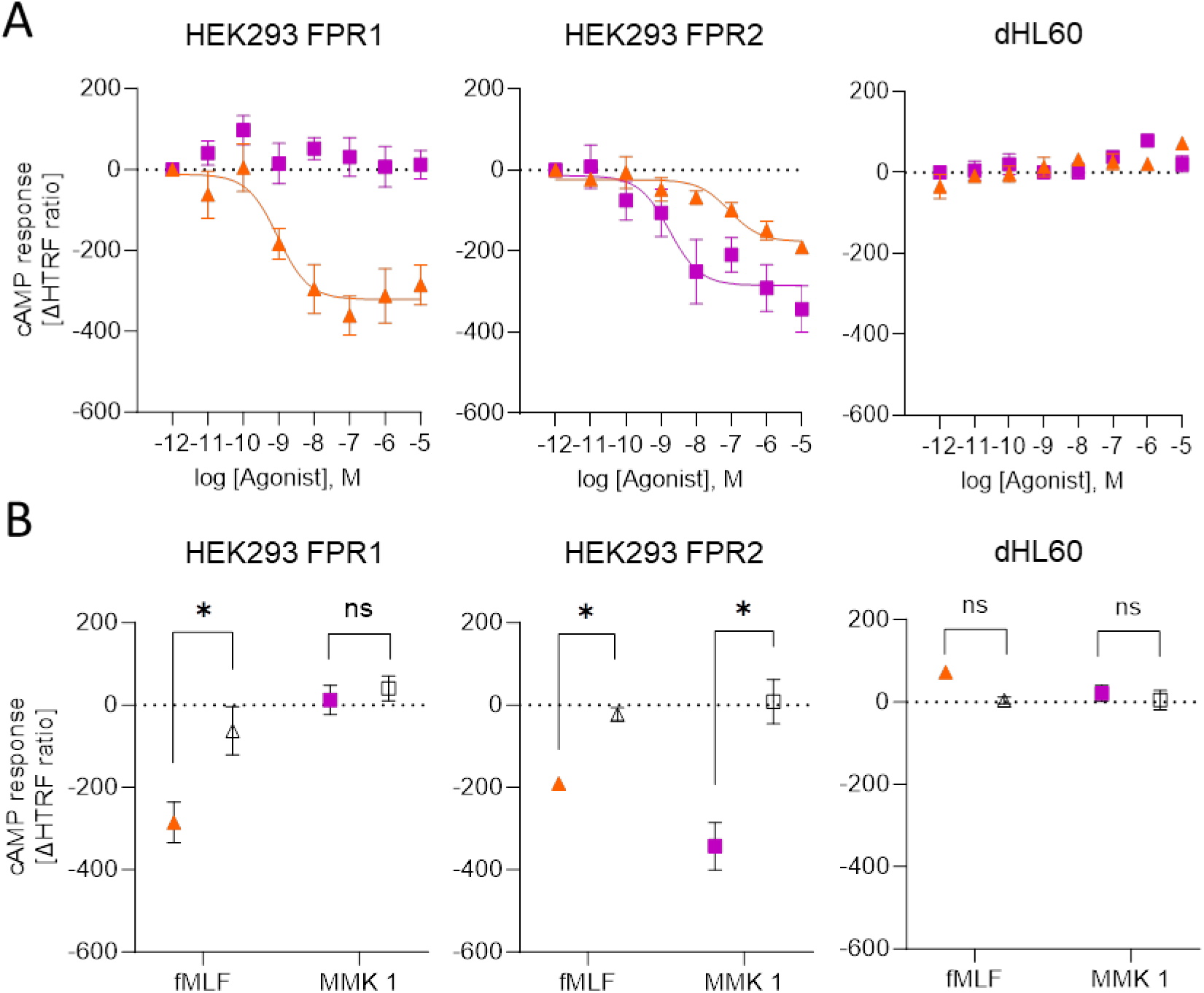
FPR-mediated changes in isoproterenol-induced cAMP formation. Concentration-dependent inhibition of isoproterenol-induced *de novo* cAMP formation was assessed for fMLF (orange triangle) and MMK 1 (magenta rectangle) in HEK293 cells stably expressing FPR1 or FPR2 as well as in dHL60. (A) Concentration-response curves were fitted using a four-parameter logistic model with the Hill slope set to 1, data points indicate the mean ΔHTRF ratios ± SEM of at least 5 independent experiments. (B) Statistical analysis of the differences in responses at the lowest and the highest agonist concentrations. Mean ΔHTRF ratios ± SEM were compared by two-tailed Students’ *t*-test, ns, not significant, *, p-value < 0.05.

### G protein-dependent phosphorylation of ERK1/2 occurs regardless of the cellular environment

Because these findings implied that the FPR signalling response was governed by the cell environment, we next explored whether this also held for activation of the ERK1/2 MAPK signalling pathway, which has been linked to these receptors [23,32]. Contrary to the cell-type-dependent differences in cAMP regulation, a concentration-dependent increase in ERK1/2 phosphorylation was detected upon stimulation with the FPR agonists in all cell lines (Fig. 3A), thus excluding that the agonists were not able to activate the naturally expressed FPRs in the dHL60 cells. ERK1/2 activation elicited by 10^-5^ M each of the agonists with or without PTX was strongly prevented by PTX pre-treatment in all systems (Fig. 3B) and confirmed Gαi-driven pathway activation [22,32]. Thus, receptor activation was indeed coupled to G protein signalling *per se*; however, specific signalling pathways were impacted by the cell environment.

**Figure 3.**
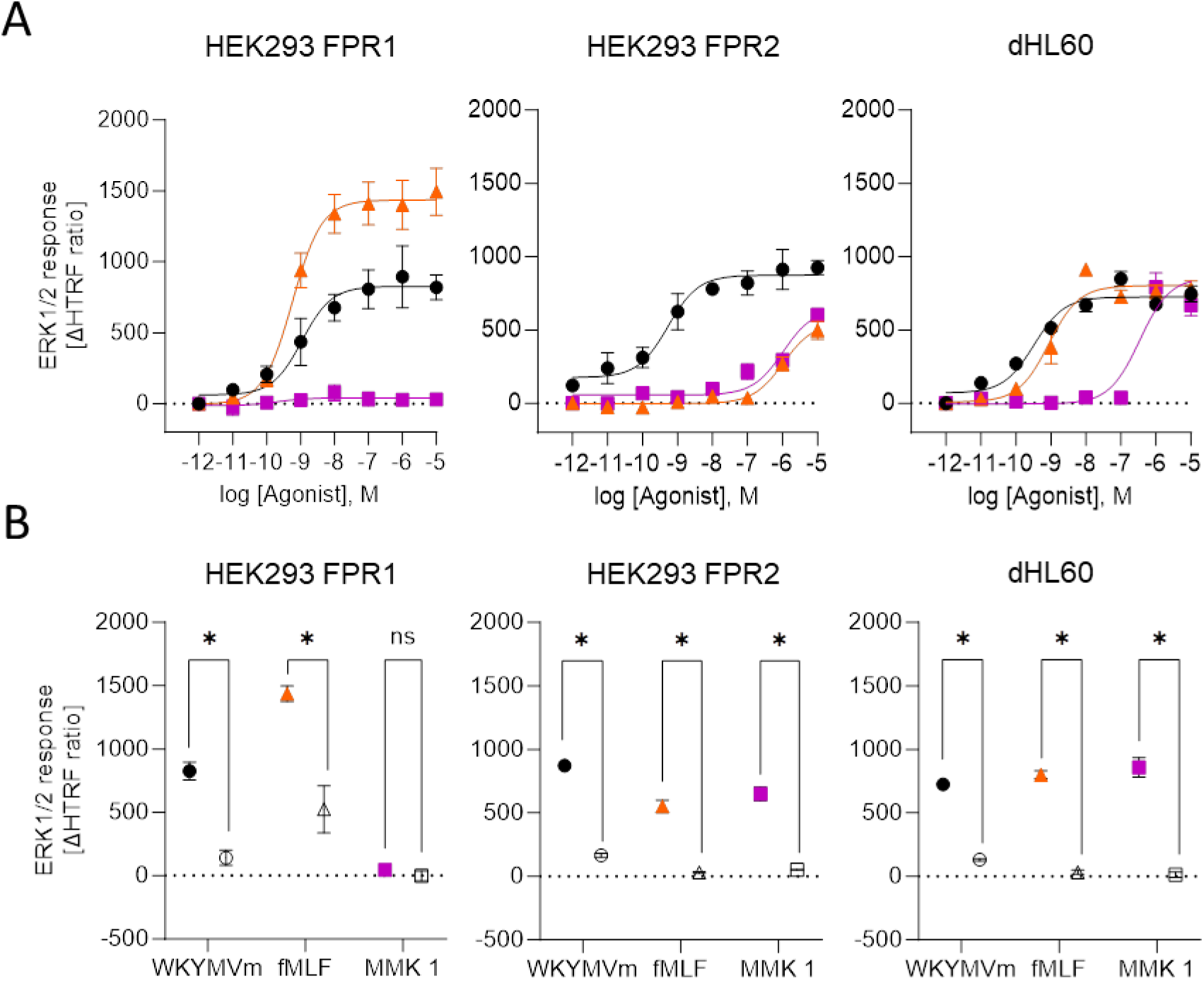
FPR-driven MAPK signalling is transmitted via Gai coupling. (A) ERK1/2 phosphorylation was assessed upon stimulation with WKYMVm (black circle), fMLF (orange triangle) or MMK 1 (magenta rectangle) in HEK293 FPR1, HEK293 FPR2 and dHL60 cells. Concentration-dependent responses were recorded 5 min after agonist addition. Concentration-response curves were fitted using a four-parameter logistic model with the Hill slope set to 1. (B) Cells were treated with 10^-5^ M each of the agonists with or without PTX. Responses were compared by a two-tailed Student’s *t*-test. Data points represent the mean ΔHTRF ratios ± SEM of at least 5 independent experiments. ns, not significant, * p-value < 0.05.

### System-specific expression of the Gαi-interacting adenylyl cyclases

While our data on PTX-sensitive ERK1/2 activation confirmed the functionality of Gai-coupling in all cell lines, our findings revealed the specific impact of the cell environment on FPR-mediated cAMP regulation. Therefore, we analyzed the expression of the adenylyl cyclases ADCY5 and 6 as the immediate downstream effectors of the Gai-containing heterotrimeric G protein complexes by qPCR. The results revealed a divergent ADCY isoform expression pattern; whereas these Gai-sensitive adenylyl cyclases were reliably detected in the HEK293-derived cell lines, their mRNA expression levels failed to achieve the prespecified minimum signal intensity in the neutrophil-like dHL60 and also in human primary PMN (Fig. 4A). As mature neutrophils exhibit low basal transcriptional activity, leading to the presence of neutrophil proteins even when their corresponding mRNA levels are low [33], we verified the presence or absence of ADCY 5 and 6 proteins by immunoblotting (Fig. 4B). Importantly, the protein expression pattern of these critical ADCYs mirrored the qPCR data. ADCY5 and 6 were found in the HEK293-derived cell lines but were neither detected in the neutrophil-like dHL60 nor PMN (Fig. 4B).

**Figure 4.**
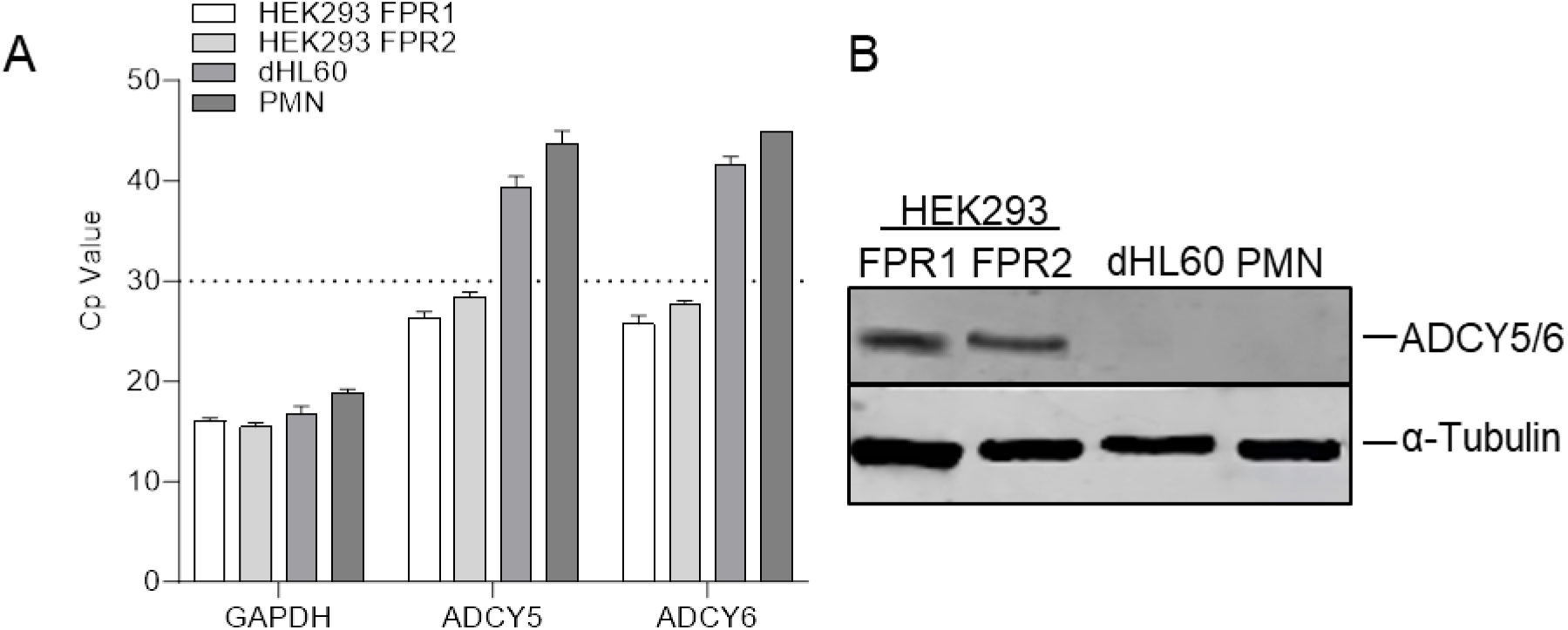
Comparison of adenylyl cyclase expression. HEK293 FPR1, HEK293 FPR2, dHL60, and PMNs were analyzed for ADCY5 and ADCY6 expression by A) qRT-PCR with GAPDH as the reference gene. Bar graphs represent the mean ± SEM of at least 3 individual experiments. A Cp value ≥ 30 (dotted line) was set as the threshold for reliable detection. B) To detect ADCY protein levels, cell lysates were analyzed by Western blot with α-tubulin as a loading control.

### Physiological bias is reflected in primary human neutrophils and also influences the classical neutrophil chemoattractant receptor BLT1

As anticipated, FPR-mediated signalling in PMN recapitulated the pattern in the neutrophil-like dHL60 cells, i.e. no significant changes in *de novo* cAMP generation (Fig. 5A, B) but efficient activation of ERK1/2 signalling (Fig. 5C). For the detection of ERK1/2 activation in PMNs, we resorted to a FACS-based assay. This approach was consistent with the results obtained by HTRF (Suppl. Fig. S1D). We, therefore, reasoned that the absence of the Gai-sensitive ADCYs would affect the impact of stimuli transmitted via Gai-coupled receptors in general.

**Figure 5.**
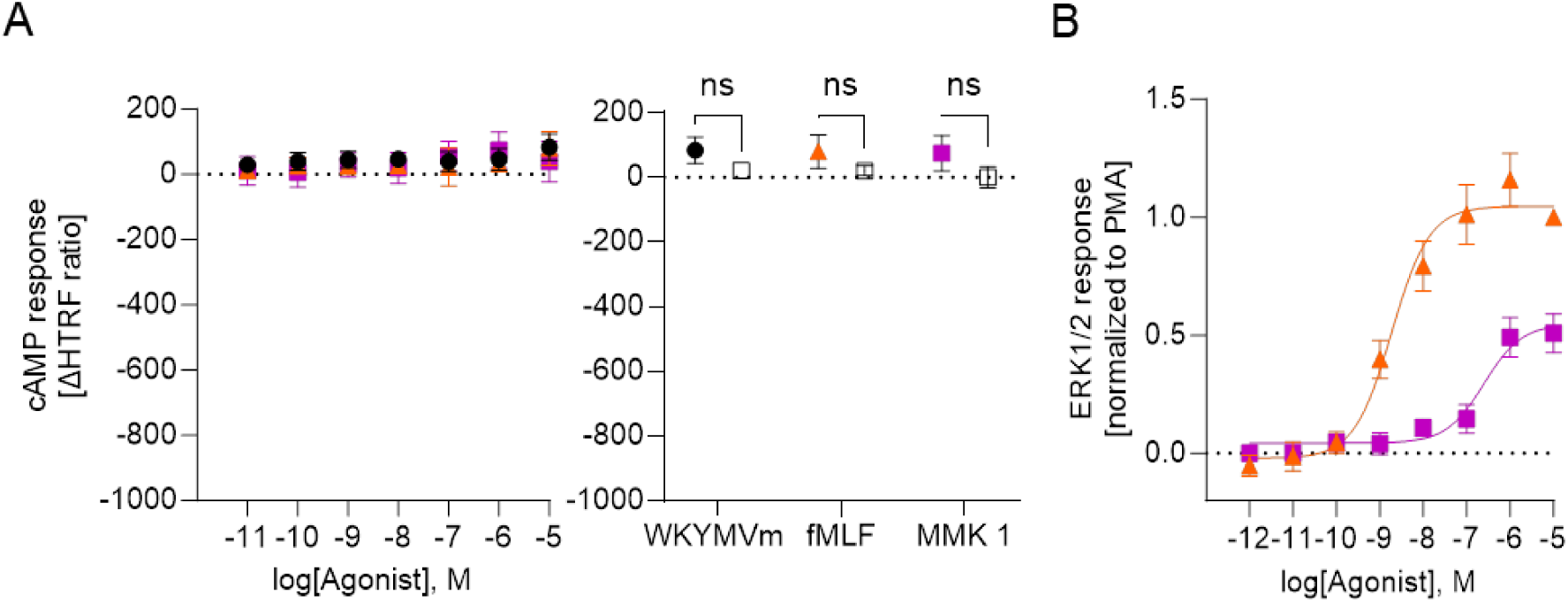
FPR-driven responses in human primary neutrophils. (A) FPR-mediated changes in *de novo* cAMP formation were assessed 30 min after agonist addition (W-peptide, black circle, fMLF, orange triangle, MMK 1, magenta rectangle) across a range of concentrations. Data points represent mean ΔHTRF ratio values ± SEM of at least 5 independent experiments. Mean ± SEM ΔHTRF ratios at the lowest and the highest agonist concentrations were compared by two-tailed Student’s-test. (B) ERK1/2 phosphorylation was assessed upon stimulation with fMLF (orange triangle) or MMK 1 (magenta rectangle) and dose-dependent responses were recorded 5 min after agonist addition. Responses were normalized to the max response obtained with PMA. To compute the logistic parameters, concentration-response curves were fitted using a four-parameter logistic model with the Hill slope set to 1. Data points represent the mean values ± SEM of at least 5 independent experiments.

To explore whether Gαi-coupled GPCRs expressed in neutrophils are incapable of regulating cAMP levels in general, we next assessed whether the LTB4 high-affinity receptor BLT1 follows a similar signalling pattern. We first probed LTB4-induced responses on *de novo* cAMP formation 30 min after agonist addition. In line with the absence of the Gai-sensitive ADCY5 and ADCY6, we did not detect an impact of LTB4 on the cAMP formation (Fig. 6A). Nevertheless, immunoblot analysis of LTB4-treated PMN showed a clear and significant ERK1/2 activation after 15 min, similar to fMLF activation (Fig. 6B), indicating that BLT1 was able to transduce signals upon specific activation.

**Figure 6.**
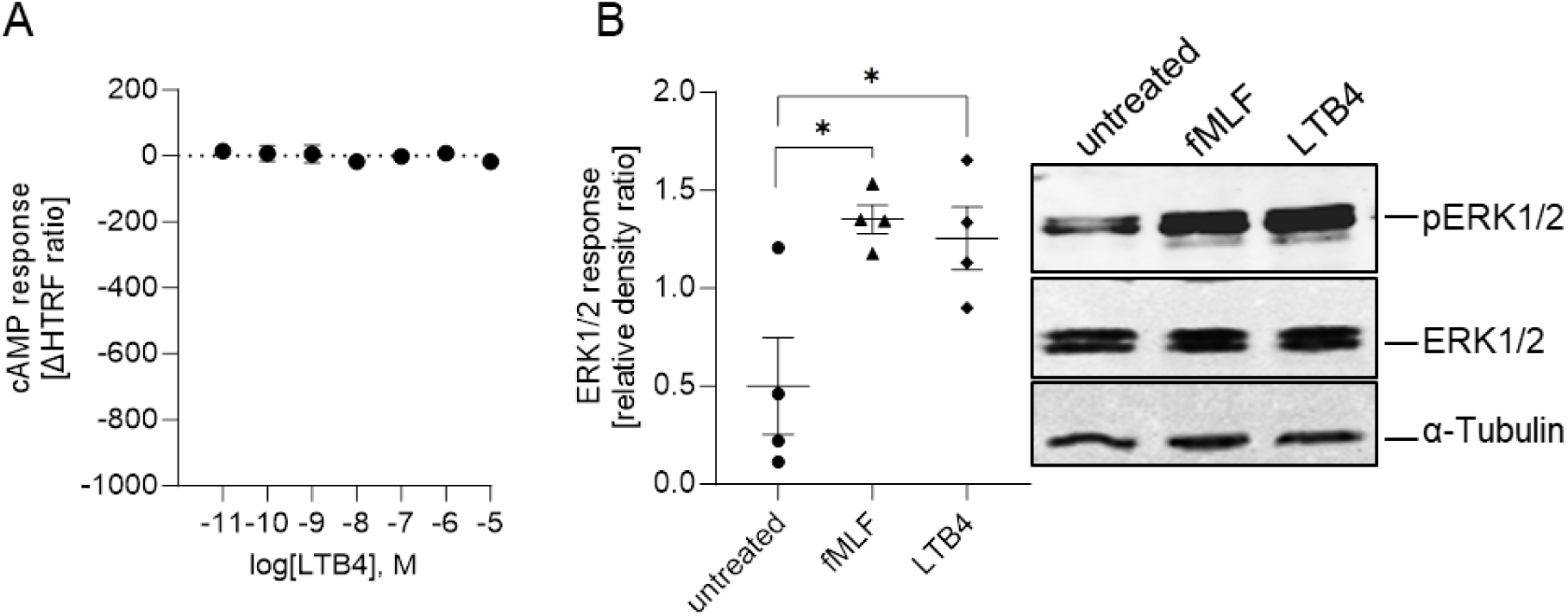
LTB4-induced responses on *de novo* cAMP formation and ERK1/2 activation in human primary neutrophils. (A) LTB4-mediated changes in *de novo* cAMP formation were assessed 30 min after agonist addition. Data points represent mean ΔHTRF ratio values ± SEM of at least 5 independent experiments. (B) PMN were treated with or without LTB4 for 15 min, with fMLF stimulation for 5 min as a positive control. After the detection of phospho-ERK1/2, the blots were stripped and probed for total ERK1/2. A representative blot of 4 independently performed experiments is shown. Tubulin was used as an additional loading control. The mean intensities of the relative phospho-ERK1/2 density ratios were compared by one-way repeated measures ANOVA and Dunnett’s post-hoc testing, * p < 0.05.

### Physiological bias is conserved across species

We also wondered whether the lack of inhibitory control on cAMP generation was a general theme seen in neutrophils across animal lineages. We therefore repeated the experiments in functional murine neutrophils differentiated *ex vivo* from HoxB8, conditionally immortalized mouse hematopoietic progenitor cells. The suitability of such neutrophils derived from the Hoxb8 progenitors to function *in vivo* in murine neutrophil-mediated inflammation models has been verified by adoptive transfer studies [34], thus underlining the relevance of this model, especially with regard to the 3R principles. Analysis of ADCY expression by qPCR revealed that the *ex vivo*-generated murine neutrophilic cells also lack ADCY 5 and 6 expressions (Fig. 7A, for the qPCR-based screen of the nine transmembrane ADCY isoforms in mice see Suppl. Fig. S3). Accordingly, the forskolin-induced cAMP formation was neither blocked by activation of the FPRs nor by LTB4-mediated BLT1 activation (Fig. 7B, C), yet significant ERK1/2 activation was observed (Fig. 7D), arguing for a conserved mechanism of G protein-dependent cAMP handling across species in neutrophils.

**Figure 7.**
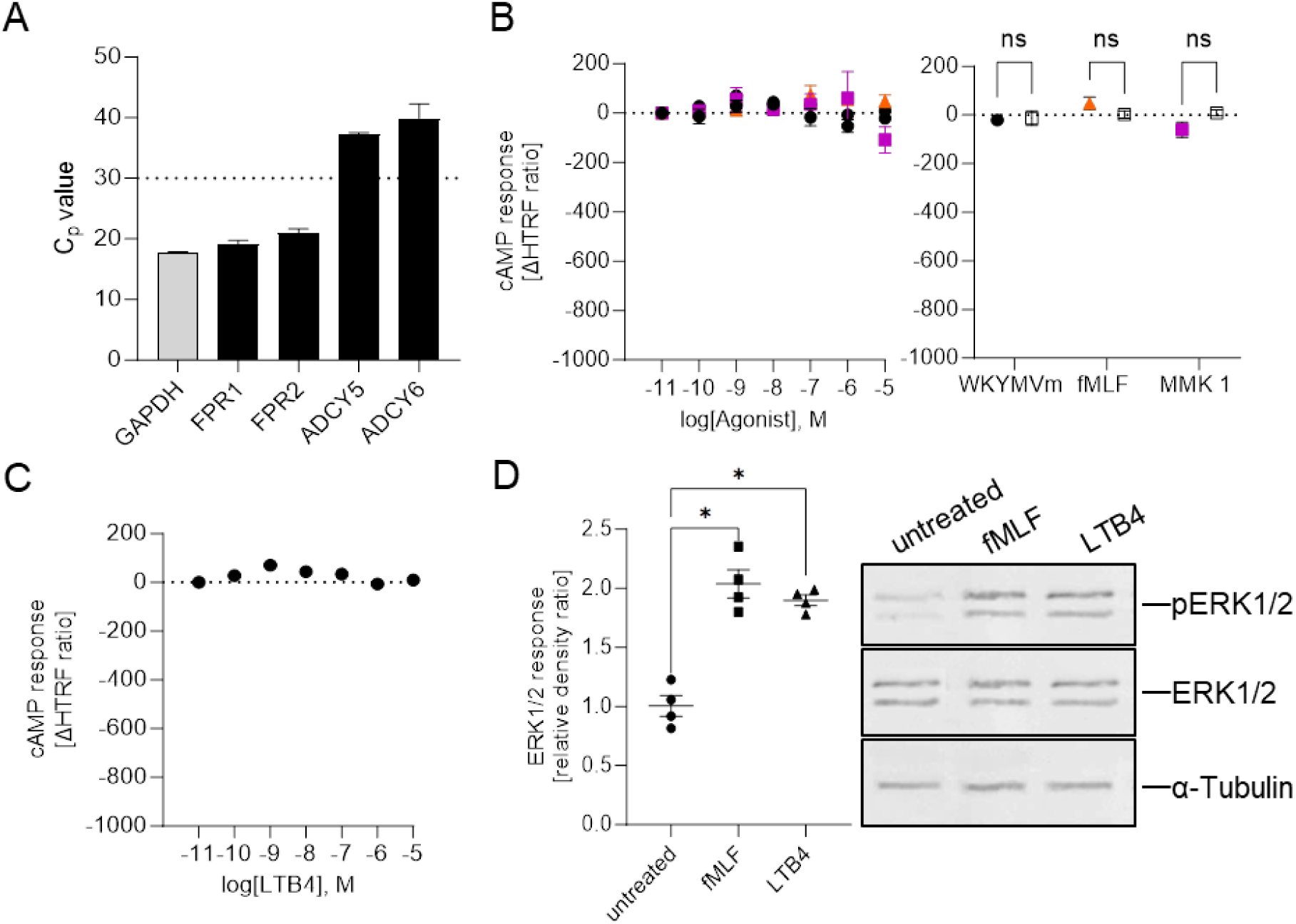
The neutrophilic Gai-coupled functional profile is conserved across species. (A) The ADCY expression profile in Hoxb8-derived neutrophils was determined by qRT-PCR. Data represent means ± SEM of Cp values of 5 replicates, with GAPDH as the reference gene. Assessment of changes in *de novo* cAMP formation upon (B) FPR (W-peptide, black circle, fMLF, orange triangle, MMK 1, magenta rectangle) and (C) LTB4 stimulation of Hoxb8 progenitor-generated neutrophils. Data points indicate the mean ΔHTRF ratios ± SEM of at least 5 independent experiments. (D) LTB4- and fMLF-induced ERK1/2 phosphorylation was assessed by immunoblotting. To detect total ERK1/2, phophoERK1/2-stained blots were stripped and reprobed. Tubulin was used as an additional loading control. A representative blot of 4 independently performed experiments is shown. The mean intensities of the relative phospho-ERK1/2 density ratios were compared by one-way repeated measures ANOVA and Dunnett’s post-hoc testing, * p < 0.05.

## Discussion

The concept of “biased signalling” explains the observations that different ligands activate specific subsets of the entire signalling repertoire of their shared receptor [14]. Initially applied to ligand-specific responses at a single receptor, this concept has evolved to encompass more intricate signal integration at higher complexity levels [35–37]. GPCR responses are also governed by non-ligand molecules such as G proteins, GPCR kinases (GRKs), arrestins, and downstream effectors and modulators which might differ in their spatiotemporal expression patterns across tissues [17]. Consequently, cell type-specific responses triggered by the same agonist-receptor pair are envisioned yet their physiological relevance remains so far unexplored.

The activation of ADCYs triggers the production of cAMP from ATP, leading to elevated intracellular levels of the second messenger cAMP [10]. GPCR signalling through Gαi typically inhibits adenylyl cyclases, thus blocking *de novo* cAMP generation by other cellular stimuli. Our comparative analysis of FPR signalling in FPR-transfected HEK293 cells, a classical model in GPCR-based drug discovery [19], and PMNs as the FPR natural environment [38–40] revealed divergent outcomes, highlighting the influence of cellular context. Our selection of receptor-specific ligands activating only the designated GPCRs ensured the unambiguous assignment of the observed responses. In neutrophils, activated GPCRs elicited Gαi-dependent signalling, however, these cells were unable to counteract Gαs-stimulated cAMP formation, whether stimulated directly or by β-adrenoceptor activation. Therefore, this cell type-dependent deviation from the textbook scheme underscores physiological bias.

Our analysis revealed the absence of adenylyl cyclases ADCY5 and ADCY6 the Gαi-inhibitable GPCR effectors [12] in neutrophils. As ADCY 5 and especially ADCY6 are prone to forskolin-induced activation [41], their absence also might explain that PMNs were rather insensitive to this diterpene. Deep-sequencing transcriptome data of neutrophil and neutrophil-like HL60 cells confirmed the expression data [42]. Importantly, this expression pattern was mirrored in *ex vivo*-differentiated murine neutrophils [34], implying that the functional bias is evolutionarily preserved. The retained ability of the stimulated receptors to activate ERK1/2, which has been demonstrated to also depend on Gi activation [22,32] affirms the functionality of the receptor-transducer axis and highlights the pivotal role of the effector proteins in physiological bias.

Beyond the Gαi-coupled FPRs and BLT1 [43–45], most chemoattractant receptors signal through pertussis toxin-sensitive pathways, indicating Gαi coupling [21,46]. The compromised downstream effectors of Gαi signalling in human and mouse neutrophils raise questions about the general ability of this cell type to regulate ADCY activation, leaving the preservation of Gαi coupling of GPCRs in neutrophils an open question beyond the scope of this manuscript.

Ligand and system bias jointly shape the functional selectivity of an agonist. Traditional heterologous and ex vivo cell systems primarily assess ligand bias on a fixed cellular background, making it challenging to discern the contribution of each factor in complex tissues or organisms. Our findings emphasize the significance of the cellular environment and provide a conclusive example of this so far only conceptually discussed physiological bias. The critical role of cellular context holds significant implications for drug development, particularly in leveraging Gαi-coupled pathways in immune cells.

## Materials and Methods

### Reagents

The cAMP inducer forskolin, the phosphodiesterase inhibitor 3-isobutyl-1-methylxanthine (IBMX), and the selective β-adrenoceptor agonist isoproterenol were purchased from Sigma Aldrich. The Gai-inhibitor pertussis toxin (PTai-inhibitor pertussis toxin (P) and the FPR agonists WKYMVm, fMLF, and MMK 1 were obtained from Tocris, the BLT1 agonist LTB4 was obtained from Cayman Chemicals.

### Cell lines and culture conditions

The HEK293 FPR cell lines stably expressing either FPR1 or FPR2 fused to a FLAG-tag were generated as previously described [25] and were cultured in Dulbecco’s modified Eagle’s medium (DMEM, PAN Biotech, Germany) supplemented with 10% standardized fetal bovine serum (FBS Advanced, Capricorn-Scientific, Germany), 100 U/mL penicillin, 0.1 mg/mL streptomycin (GE Healthcare, USA), 1% L-glutamine, 1% non-essential amino acids (NEAA, Sigma) at 37° C and 7% CO_2_.

The human promyelocyte cell line HL60 was cultured in Roswell Park Memorial Institute (RPMI) 1640 very low endotoxin medium (RPMI-VLE, PAN Biotech, Germany) supplemented with 10% endotoxin-free fetal calf serum (FCS Gold, PAA), 100 U/mL penicillin, 0.1 mg/mL streptomycin (GE Healthcare, USA), 1% L-glutamine, 1% non-essential amino acids (NEAA, Sigma) at 37° C and 5% CO_2_. To induce differentiation toward neutrophil-like dHL60 cells, 1.25% DMSO was added to the medium for 6 days.

Murine neutrophils were generated *ex vivo* from immortalized neutrophil progenitors expressing the Hoxb8 transcription factor fused to the human estrogen receptor (Hoxb8-ER) essentially as described [34]. The immortalized Hoxb8 cells were cultured in OptiMEM Glutamax medium (Fisher Scientific) supplemented with 100 U/mL penicillin, 0.1 mg/mL streptomycin, 30 µM beta-mercaptoethanol, 10% fetal bovine serum (FBS) (Biochrome, Cambridge, UK), 20 ng/mL recombinant SCF (ImmunoTools, Friesoythe, Germany), and 1 µM β-estradiol at 37°C and 5% CO2. To induce neutrophil differentiation, cells were transferred into an estrogen-free medium for 4 days. Assessment of the complete downregulation of c-Kit and CD34, and upregulation of CD11b and Ly6G verified the successful establishment of the neutrophil phenotype [47,48].

All cell lines were routinely tested for the absence of mycoplasma contamination.

### Isolation of human peripheral blood neutrophils

Human primary polymorphonuclear neutrophils (PMN) were isolated from peripheral blood collected from healthy volunteers upon informed consent using polymorphprep™ from Progen according to the manufacturer’s protocol and as described before [25]. Briefly, 1 volume of blood was added to 1.5 volumes of pre-warmed polymorphprep. After centrifugation for 35 min at 500 x g (no breaks) at room temperature (RT), the PMN phase was collected and washed with PBS. Following erythrocytes were lysed by incubation with 1xRBC Lysis buffer (BioLegend), PMNs were resuspended and stored in Hanks balanced salt solution (HBSS) with Ca^2+^, and Mg^2+^, supplemented with 10 mM HEPES, 0.1% glucose, and 0.25% BSA, pH 7.4. Blood drawing was approved by the institutional review board.

### Quantitative reverse transcription PCR

Gene expression levels of the FPRs and ADCY isotypes were analyzed via RT-qPCR. RNA isolation was performed with the RNeasy mini kit (Qiagen, Germany) according to the manufacturer’s protocol, and 1 µg RNA starting material was reverse-transcribed with the High-Capacity cDNA Reverse Transcription Kit (Thermo Fisher Scientific, Germany). Gene expression was analyzed with predesigned QuantiTect primer assays (Qiagen) or custom-designed Primer (Suppl. Table 1). The qPCR reactions were performed with the Brilliant III Ultra-Fast SYBR Green qPCR Master Mix (Agilent Technologies, USA) on the Roche LightCycler480 PCR system (1 min 95 °C; 50 cycles of 5 sec at 95 °C, 10 sec 60 °C and final melting curve at 95 °C).

### Evaluation of adenylyl cyclase IV/V/VI protein expression by immunoblotting

ADCY V/V/ protein levels were assessed by immunoblotting. Cells were lysed with RIPA buffer (25 mM Tris-HCl pH7.6, 150 mM NaCl, 1% NP-40, 1% sodium deoxycholate, 0.1%SDS; supplemented with protease and phosphatase inhibitors). Proteins were resolved by SDS-PAGE and transferred onto nitrocellulose membrane (GE Healthcare, Amersham™ Protan™ 0.2µM NC). ADCY bands were immunostained with mouse anti-ADCY5/6 (Santa Cruz; diluted 1:400 in blocking buffer), and mouse-anti-α -tubulin (Sigma Aldrich, 1:2000 dilution in Intercept Blocking Buffer). IR-Dye 680RD goat-anti-mouse and IR-Dye 800RW donkey-anti-rabbit antibodies (Li-COR, 1:10000 dilution in Intercept Blocking Buffer) were used as the secondary antibodies. Bands were visualized on the Odyssey M Imaging System (LI-COR).

### HTRF-based quantification of cAMP levels

FPR agonist-induced changes in the cellular *de novo* cAMP formation were measured with the HTRF-based competitive cAMP-Gi Kit (Cisbio, Revity) as previously published [23]. In brief, cells cultured in 96 well plates (50000 cells/well) were starved for 30 min in DMEM or RPMI supplemented with 500 µM IBMX. Cells were either activated with 5 µM forskolin or left untreated. All cell lines were further incubated with FPR agonists at indicated concentrations for 30 min at 37 °C. Cells were lysed and transferred to a 384-well plate (Greiner Bio-one), conjugates were added, and samples were incubated for 1 hour at RT in the dark. The CLARIOstar reader (BMG Labtech) (200 flashes/well, integration start 60 µsec, integration time 400 µsec, settling time 100 µsec) was utilized to measure luminescence signals that were expressed as the ratio of 10.000 x (620/665nm). Initial baselines were subtracted from the ratios recorded upon stimulation to calculate ΔHTRF values.

### Analysis of MAPK/ERK pathway activation

ERK1/2 phosphorylation in HEK293 cells was measured with the HTRF (Homogeneous Time-Resolved Fluorescence) immunoassay from Cisbio, as described previously [23]. In brief, cells were seeded in a 96-well plate (30000cells/well/25µL) in serum-free DMEM followed by stimulation with selected agonists at the indicated concentrations for 5 min at 37 °C. Subsequently, cells were lysed with supplemented lysis buffer for 30 min and then transferred to a 384-well plate. Labeled conjugates were added and the plate was incubated for 4 hours at RT in the dark. The luminescence was recorded with the CLARIOstar reader (BMG Labtech) and the signals were expressed as the ratio of 10000 x (acceptor signal/donor signal). Initial baselines were subtracted from the ratios recorded upon stimulation to calculate ΔHTRF values. FPR-mediated ERK1/2 phosphorylation in dHL60 and primary neutrophils was measured via flow cytometry. In brief, 2×10^5^ cells were incubated with agonists at indicated concentrations for 5 min at 37 °C, followed by fixation with 4% PFA. Permeabilization was performed at −20 °C for an hour with the True-Phos™ Perm Buffer from Biolegend. Staining was performed with FITC-coupled mouse monoclonal anti-ERK1/2 phospho (Thr202/Tyr204, Biolegend) antibody (diluted 1:200 in staining buffer) for 30 min at RT in the dark. The number of positive cells was recorded on a Guava EasyCyte flow cytometer (Millipore). To assess BLT-1-mediated ERK activation in primary neutrophils and Hoxb8-derived murine neutrophils, 1-3×10^6^ cells were treated with LTB4 (1µM) or the respective controls for 15 min and were then lysed in RIPA buffer (25 mM Tris-HCl pH7.6, 150mMNaCl, 1% NP-40, 1% sodium deoxycholate, 0.1%SDS; supplemented with protease and phosphatase inhibitors). Proteins were resolved by SDS-PAGE and transferred onto nitrocellulose membrane (GE Healthcare, Amersham™ Protan™ 0.2µM NC). After detection of phospho-ERK1/2 by immunostaining with mouse anti-P-p44/42 MAPK XP (Cell Signalling, diluted 1:500 in Intercept Blocking buffer, Li-COR), the blots were stripped and probed for total ERK1/2 (rabbit anti-p44/42 MAPK antibodies, Cell Signalling, diluted 1:500 in Intercept Blocking buffer, Li-COR), and mouse-anti-alpha-tubulin as loading control (Sigma Aldrich, 1:1,000 dilution in Intercept Blocking Buffer). IR-Dye 680RD goat-anti-mouse and IR-Dye 800RW donkey-anti-rabbit antibodies (Li-COR, 1:10000 dilution in Intercept Blocking Buffer) were used as the secondary antibody and were visualized on the Odyssey M Imaging System (LI-COR). Signal intensities were quantified from the profile plot of each lane using ImageJ (Rasband, W.S., ImageJ, U. S. National Institutes of Health, Bethesda, Maryland, USA, https://imagej.net/ij/, 1997-2018) and the relative densities were calculated as described in [49].

### Statistical Analysis

Concentration-response curves were generated utilizing a four-parameter logistic model with the Hill slope set to 1 and logistic parameters were extrapolated (GraphPad Prism 9 software). Outliers were detected by ROUT testing with a Q-value threshold set to 1%. Mean values ± standard error of the mean (SEM) were calculated from independent experiments as indicated and were assessed for statistically significant differences by two-tailed Student’s t-test (two groups) or one-way repeated measures ANOVA and Dunnett’s post-hoc testing. A p-value < 0.05 was set as the threshold for statistical significance.

## Author Contributions

DP, LP, CP and MFS performed the experiments and collected and analyzed the data. OF, MFS, OS, and TV provided human and murine neutrophils and further resources. MB, DP, CAR, and UR designed and supervised the research, analyzed the data and wrote the paper.

## Acknowledgement

This work is funded by the Deutsche Forschungsgemeinschaft (DFG, German Research Foundation) – CRC1009 “Breaking Barriers” project A06 to UR. and supported by the Graduate School of Natural Products (GS-NP).

## Supporting Information

**Suppl. Table S 1:**
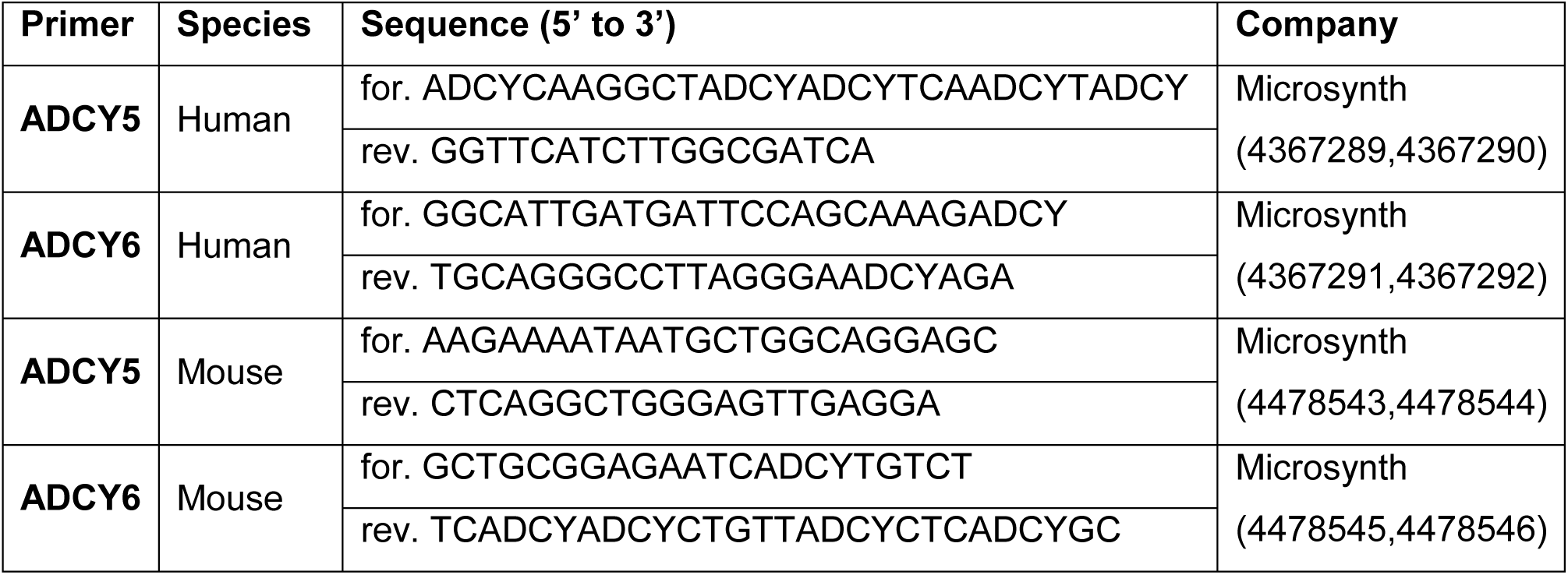
Primer pairs of ADCY5 and ADCY6 for the detection of gene expression in human and mouse.

**Suppl. Figure S1:**
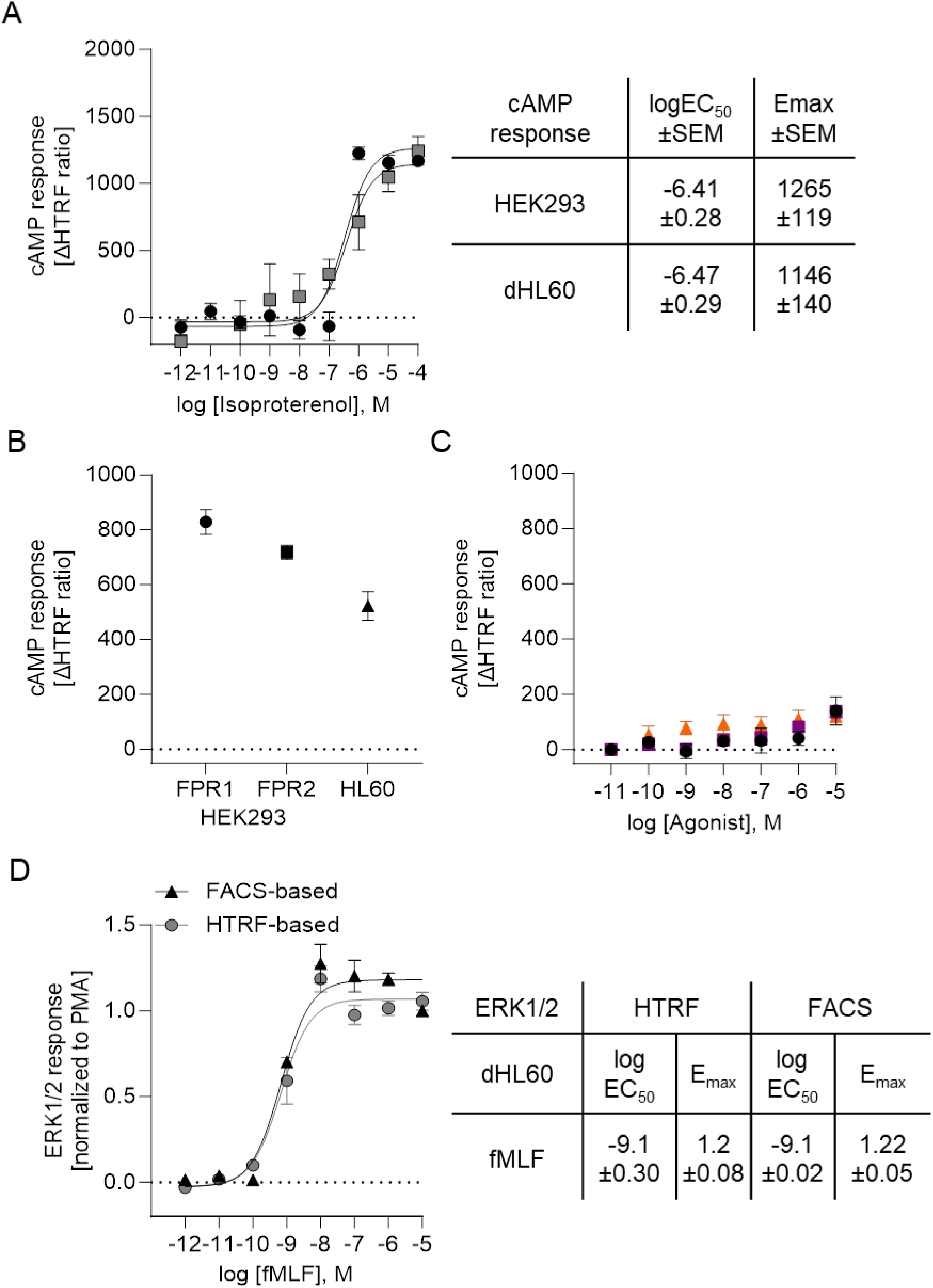
System validation. (A) HEK293 cells (grey squares) and dHL60 cells (black circles) were stimulated with isoproterenol and ΔHTRF ratios were recorded 30 min after agonist addition. Data points represent the mean ± SEM ΔHTRF ratios of at least 5 independent experiments. Concentration-response curves were fitted using a four-parameter logistic model with the Hill slope set to 1. Logistic parameters ±SEM are compared in the table. (B) FSK-induced changed in de novo cAMP generation in the HEK293 and dHL60 cell lines. Data points represent the mean ± SEM ΔHTRF ratios of at least 5 independent experiments. (C) dHL60 cells were stimulated with the FPR agonists WKYMVm (black circle), fMLF (orange triangle), and MMK 1 (magenta rectangle). ΔHTRF ratios were recorded 30 min after agonist addition. Data points represent the mean ± SEM ΔHTRF ratios of at least 5 independent experiments (D) dHL60 cells were stimulated with fMLF. Concentration-dependent ERK1/2 activation was quantified by flow cytometry (grey circles) and HTRF black triangles). Responses were recorded 5 min after agonist addition and were normalized to the PMA response. Concentration-response curves were fitted using a four-parameter logistic model with the Hill slope set to 1. Data points represent the response ±SEM of 3 experiments. Logistic parameters are compared in the table

**Suppl. Figure S2:**
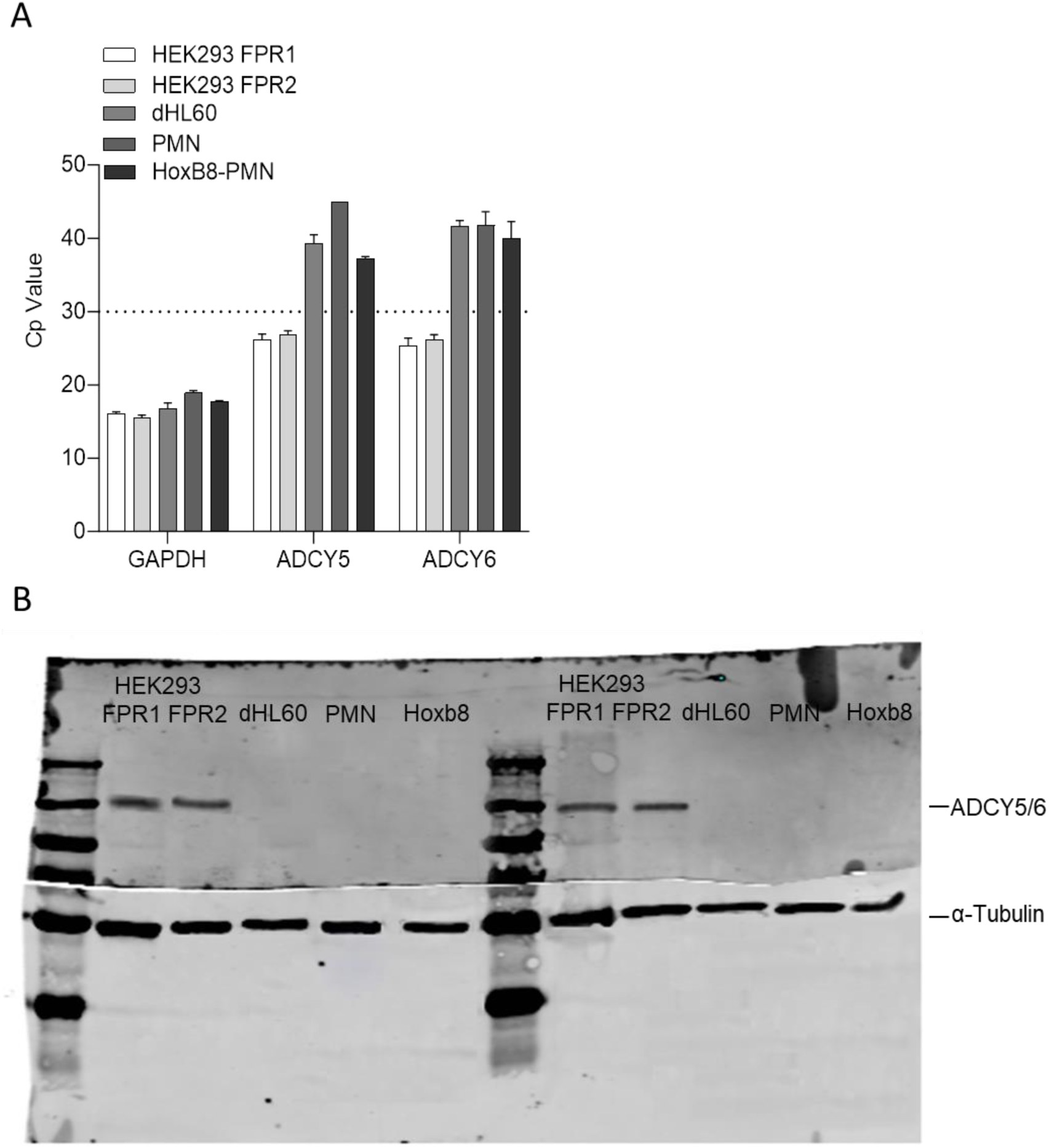
Expression analysis of human and mouse transmembrane adenylyl cyclase isoforms V/VI. A) ADCY V/VI expression in HEK293 FPR1, HEK293 FPR2, dHL60, human PMNs and murine Hoxb8-derived PMNs was determined by qRT-PCR with GAPDH as the reference gene. Bar graphs represent the mean ± SEM of at least 4 individual experiments. A Cp value ≥ 30 (dotted line) was set as the threshold for reliable detection. B) Unedited representative immunoblot verified qPCR analysis of ADCY V/VI expression at protein level in indicated cell lines, including murine Hoxb8 neutrophils. Alpha-tubulin served as the internal loading control.

